# mRNAspect: Real-Time Quality Control of Therapeutic mRNA with *N*^1^-Methylpseudouridine Bias Correction Using Nanopore Direct RNA Sequencing

**DOI:** 10.64898/2026.11.24.768843

**Authors:** Sua Lee

## Abstract

Quality attributes of therapeutic mRNA include sequence identity, transcript completeness, and poly(A) tail length. While these attributes require monitoring from development through manufacturing, each needs a separate assay. Nanopore direct RNA sequencing enables rapid quality control because it measures all three on the same molecules within a single run, and the resulting data are available during sequencing. However, *N*^1^-methylpseudouridine (m1Ψ), which often replaces uridine in therapeutic mRNAs, alters the nanopore signal, biasing basecalling and poly(A) estimation.

We present mRNAspect, a pipeline combining real-time analysis and m1Ψ bias correction. mRNAspect includes a fine-tuned basecaller and a poly(A) correction based on the reference sequence. mRNAspect is a packaged macOS app that ingests POD5 or BAM files by watching the MinKNOW output directory as sequencing proceeds. Its graphical user interface shows the latest snapshot of the results, and it writes HTML reports as each input file is processed and a PDF report when the run is finished. When measured on Linux with a single NVIDIA RTX 2080 Ti, each 3-min batch of sequencing (about 5,700 reads) was processed in 38 s.

The basecaller was fine-tuned on *in vitro* transcribed (IVT) constructs and cDNA-IVT libraries. Across 10 m1Ψ constructs held out from training, the fine-tuned basecaller reduced the error rate from 20.1% to 7.2%, compared with 3.0% for unmodified RNA. A poly(A) correction was fitted to constructs with both modified and unmodified pairs. The correction accounts for the fraction of uridine in the reference sequence and the number of uridines within the 12-nt sequence adjacent to the tail. Across eight constructs, the correction reduced the poly(A) tail length difference between modified and unmodified pairs from up to 9.0 nt to within 1.8 nt.

## 1. Introduction

mRNA therapeutics are emerging as a drug platform (Żak and Zangi, 2025). Quality attributes of therapeutic mRNA include sequence identity, transcript completeness, and poly(A) tail length (World Health Organization, 2022; United States Pharmacopeia, 2024) and must be monitored from development through manufacturing since they can change during synthesis and storage. Therefore, robust and rapid quality control (QC) is crucial for mRNA drugs. However, current QC typically relies on separate assays for each attribute, some of which are time-consuming or labor-intensive (Camperi et al., 2025).

Nanopore sequencing from Oxford Nanopore Technologies measures many mRNA attributes in a single run and therefore is a promising platform for efficient therapeutic mRNA QC. While a single nanopore cDNA sequencing assay can measure several mRNA quality attributes (Gunter et al., 2023), nanopore direct RNA sequencing (DRS) (Garalde et al., 2018) is another option for confirming mRNA identity (World Health Organization, 2022).

Another study has shown the potential of DRS for QC, measuring full-length sequence, integrity, poly(A) tail length, and 5′ capping efficiency (Chatla et al., 2026). While the DRS library preparation takes approximately 160 minutes (Oxford Nanopore Technologies, 2026c), DRS can be time-consuming (Camperi et al., 2025). Analysis of nanopore signal during sequencing is possible (Loose et al., 2016), including in DRS runs (Chang, 2019; Sneddon et al., 2024), but, to our knowledge, no publicly available tool reports construct-level QC of mRNA during the run. Analyzing DRS data by ingesting the output files written at short intervals (e.g., 180 s) throughout the run will allow assessment before sequencing completion and shorten the time needed for QC.

Therapeutic mRNAs are often made with modified nucleosides, which suppress immunostimulation (Karikó et al., 2005). While pseudouridine incorporation in mRNA has been shown to increase translation (Karikó et al., 2008), *N*^1^-methylpseudouridine (m1Ψ) is more effective at increasing protein expression and further reducing immunogenicity (Andries et al., 2015). Today, m1Ψ is known as a capable substitute for uridine (U) in therapeutic mRNA and has shown its effectiveness through COVID-19 vaccines (Nance and Meier, 2021).

RNA modification leads to biased measurements in DRS. m1Ψ raised the error rate in RNA002 chemistry (Fleming and Burrows, 2023; Gunter et al., 2023) and also in RNA004 chemistry (Kang et al., 2026), since it alters the ionic current (Kang et al., 2026). m1Ψ induces miscalling, typically U-to-C miscalling as a signature (Fleming and Burrows, 2023). We observed that the poly(A) tail of m1Ψ-modified RNA is underestimated. Since both sequence identity and the poly(A) tail are crucial attributes of mRNA, correcting m1Ψ-induced biases is essential for accurate QC. While prior work retrained the basecaller with m1Ψ-modified RNA and recovered the alignment on the same construct, it was done on RNA002 chemistry, and its RNA004 models were only for tRNA-derived RNAs (Wang et al., 2025); to our knowledge, no publicly available RNA004 basecaller has been trained on m1Ψ-modified RNA. Analysis of poly(A) tails of COVID-19 vaccines was done by DRS (Krawczyk et al., 2025), but no calibration of the poly(A) tail for m1Ψ-modified RNA compared to unmodified RNA was applied.

Here, we present mRNAspect, a real-time pipeline that measures multiple attributes such as sequence identity, transcript completeness, and poly(A) tail length within a single DRS run. mRNAspect is distributed as a packaged macOS app and provides up-to-date results during the sequencing run, integrating rapid QC and bias correction into a single workflow. The latest measurements, at both the run and construct levels, are shown in a graphical user interface (GUI) and in reports produced. Measuring the time from basecalling to report generation for raw signal files for three minutes of sequencing showed that real-time analysis is feasible. mRNAspect includes an RNA004 basecaller fine-tuned for m1Ψ, which will be made openly available, and a reference-based correction for underestimated poly(A) tails. We show that the fine-tuned basecaller reduces per-base error and read loss during alignment of m1Ψ-modified RNA, and that the poly(A) correction reduces underestimation and narrows the poly(A) tail length difference between modified and unmodified pairs.

## 2. Implementation and methods

### 2.1 Overview and architecture

mRNAspect has three modes (Table 1). When the live mode is selected, the real-time engine watches the directory for finished POD5 or BAM files by polling and by checking that each file’s size has stopped changing. Completed files are queued and processed one at a time. Skipping of queued files is off by default, but the number of files to be skipped can be specified in the configuration file to keep the analysis in real time when basecalling is slow.

**Table 1.** The three modes of mRNAspect. Users select a mode according to their situation; the table gives the input and output of each mode.

| **User's situation** | **Mode** | **Input** | **What they get** |
|---|---|---|---|
| A run is in progress | Live | MinKNOW output directory of POD5 or BAM | GUI and HTML report updated per file; final HTML + PDF |
| A finished run, raw signal | POD5 | Directory of POD5 files | Same GUI and reports, without waiting |
| Already basecalled | BAM | BAM files | Same GUI and reports, no GPU needed |

A simplified schematic of mRNAspect is shown in Fig. 1a. While several parameters for basecalling, alignment, and QC metrics can be tuned in the configuration file, options that usually differ between QC runs, such as the input and output directories and the reference sequence, can be selected in the start menu. After basecalling and alignment, the resulting BAM file is processed read by read. Each read is passed to every QC module, which updates its values. Snapshots of the current values are integrated and reported to the user through the GUI. Whenever the snapshot is rewritten, it is saved as a JSON file and as an HTML report, and the GUI refreshes and therefore reflects the latest results. A run is considered finished when MinKNOW writes final_summary_*.txt and all queued files have been processed, or when the user stops the run. Once the run is finished, the final reports are generated in both HTML and PDF formats.

**Fig. 1.**
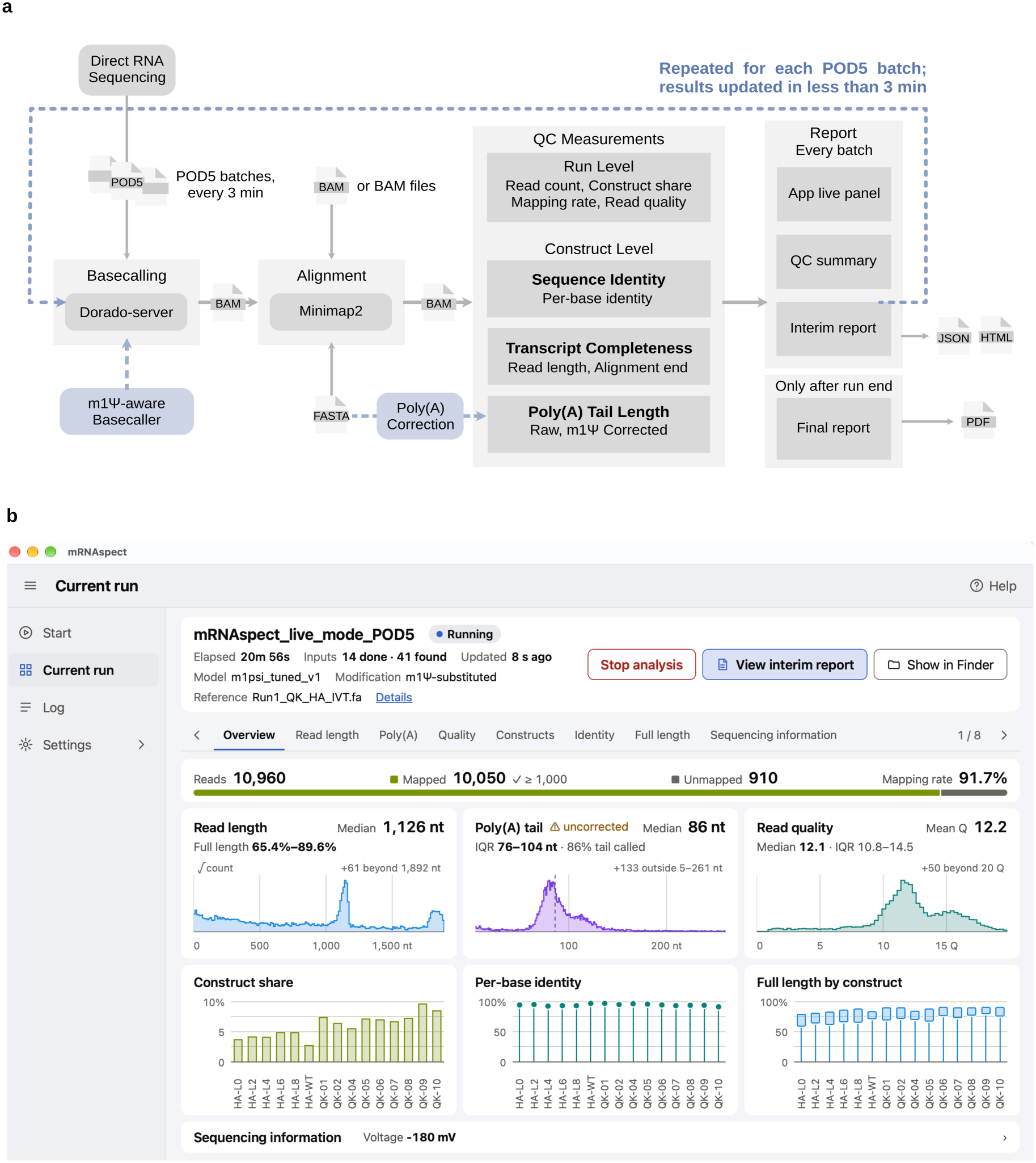
Overview of mRNAspect. **(a)** Schematic of mRNAspect. Arrows indicate the data flow, and each rectangle represents a step of the pipeline. The dashed loop represents the real-time update of the GUI and outputs during the run. Inputs and outputs are drawn as file icons. **(b)** Screenshot of the mRNAspect GUI in live mode ingesting POD5 files, showing the Overview panel of the Current-run menu. The screenshot was taken on macOS, with a sequencing run simulated by MinKNOW playback with `batch_duration = “30s”`. The demo dataset, with its reference files and configuration file, was used for the screenshot. Because the Mac used for this screenshot does not have a suitable GPU, the results shown lag behind the sequencing run.

### 2.2 Basecalling and alignment

For basecalling and alignment, the Dorado basecall server (v7.13.6, Dorado core 1.3.2; Oxford Nanopore Technologies, 2026d) and minimap2 (v2.30, r1287) (Li, 2018, 2021) are used. Basecalling is done with the `--disable_qscore_filtering` option, because m1Ψ reads could otherwise be removed for their low quality scores. The estimated poly(A) tail length is recorded in the pt tag during basecalling and then corrected for each construct. Any basecaller model can be selected, but `m1psi_tuned_v1` (hereafter the fine-tuned model) gives more interpretable results for m1Ψ-modified samples by producing fewer basecalling errors. Poly(A) tail correction is supported only for the fine-tuned model and Dorado’s rna004_130bps_hac@v5.3.0 (hereafter the stock model). After basecalling, BAM files are aligned against the user-provided reference with the default parameters `-ax map-ont -k7 -w5 --secondary=no -y`. The aligned BAM files, with the basecaller’s tags preserved, are then processed for QC.

### 2.3 QC metrics

Table 2 lists the scope, definition, and reads used for each QC metric. Metrics span run-level and construct-level measurements. While run-level measurements of read length, read quality, and mapping rate are calculated including unmapped reads, construct-level measurements and mapping quality are calculated with the reads that are assigned to a specific construct.

**Table 2.**
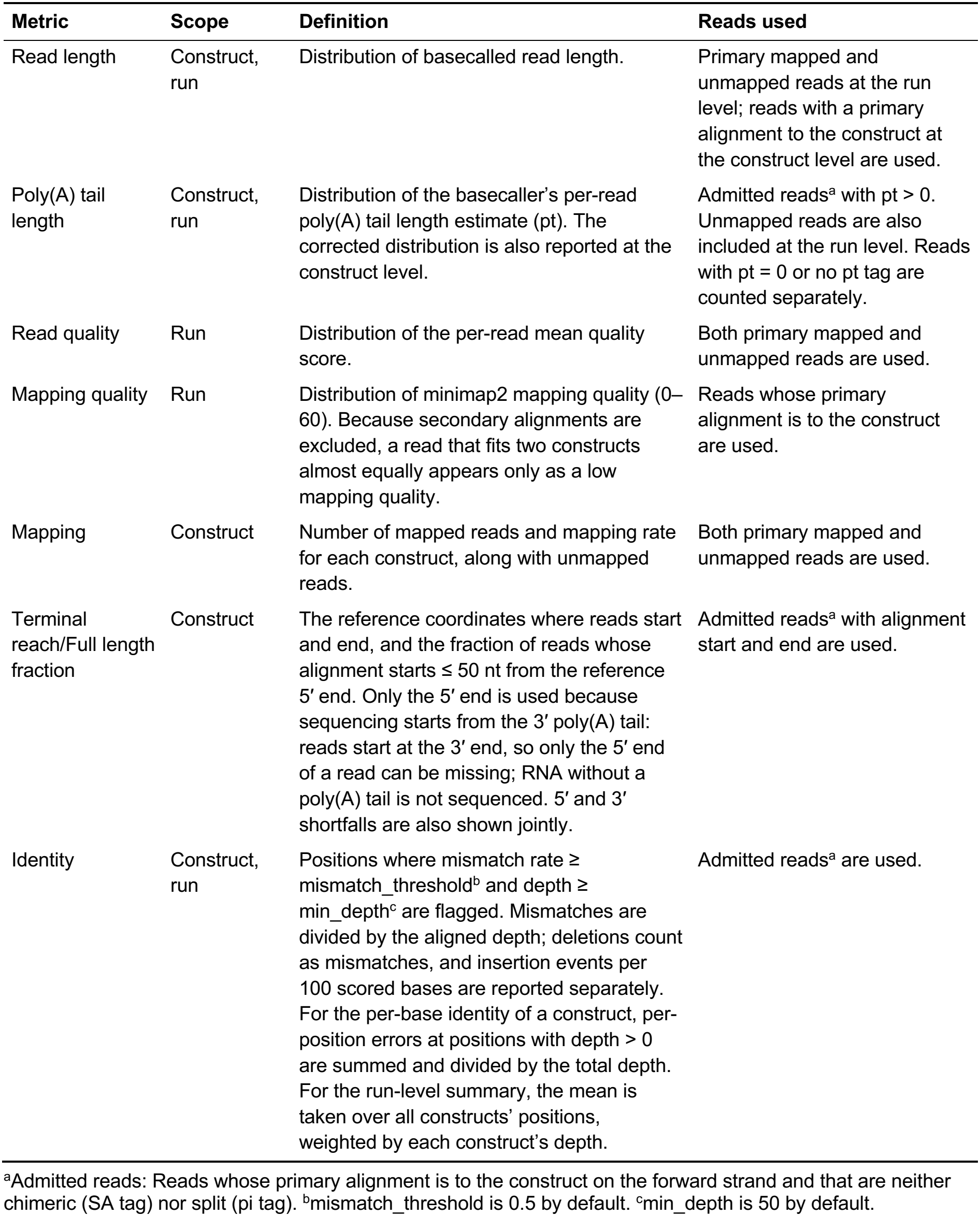
QC metrics that mRNAspect measures.

m1Ψ-modified RNAs have more signal samples per base than unmodified RNAs. Estimation of the poly(A) tail through Dorado is done by converting the poly(A) signal duration into nucleotides using the body’s samples per base (Oxford Nanopore Technologies, 2026b).

Thus, a larger number of samples per base in the body shortens the estimate, leading to underestimation of the poly(A) tail. To address this, estimated poly(A) tail length is further corrected at the construct level. The correction formula `pt_c = (pt + 10.88) × F − 10.88 − E` was fitted with the paired constructs of Run 2 (see Section 2.5). 10.88 is the offset used in Dorado estimation. F and E depend on the modification status of the RNA (unmodified or m1Ψ-modified) and on the basecaller model (stock or fine-tuned). F accounts for the U fraction of the reference sequence. For m1Ψ-modified RNA, E is computed from the number of m1Ψ in the 12 nt adjacent to the poly(A) tail. Thus, the reference sequence should end immediately before the poly(A) tail for the correction to be accurate. The fitted constructs span U fractions of 0.055–0.249 and three 3′-end sequences with 2, 3, and 6 m1Ψ in the 12 nt adjacent to the tail; a reference sequence outside these ranges is flagged. For m1Ψ-modified RNA basecalled with the stock model, alignment should be done with `-k7 -w5 --secondary=no` because the correction also accounts for the number of deleted and inserted bases.

Sequencing information is analyzed from MinKNOW output files when present. This includes end reason (the reason that a read ends; the fraction of ejected mid-molecule reads is reported separately), pore activity (the share of channel time spent in each channel state, and the strand fraction), and bias voltage over the run. If MinKNOW files for the analysis are absent, Sequencing information is not reported.

### 2.4 Basecaller fine-tuning for m1Ψ-modified RNA

The fine-tuned model was trained using Bonito v1.1.0 (Oxford Nanopore Technologies, 2026a), with --pretrained rna004_130bps_hac@v5.3.0. Fifteen m1Ψ-modified constructs and 14 unmodified constructs from two DRS runs, together with the public HeLa- and HEK293T-derived cDNA *in vitro* transcribed (cDNA-IVT) dataset (Zou et al., 2025; European Nucleotide Archive (ENA) PRJEB80229) were used in training. Evaluation was done on 10 constructs that were held out from training and validation; additionally, flank sequences shared with the training constructs were excluded. A region was excluded when a 21-nt stretch covering it also occurs in a training or validation construct (8.6% of covered positions), and the remaining positions are called novel sites. The learning rate and number of epochs were chosen from the same evaluation. Curlcake 4 (Liu et al., 2019) from public RNA004 PromethION data (Cruciani et al., 2025; ENA PRJEB82528, hereafter public Curlcake dataset), which was used in neither training nor model selection, serves as the independent test.

### 2.5 Datasets

All constructs for the DRS runs were generated by *in vitro* transcription with either UTP or m1ΨTP. Two DRS runs (Run 1 and Run 2) were performed with SQK-RNA004 on a MinION sequencer with FLO-MIN004RA flow cells. Run 1 was not multiplexed: all constructs carried the same WarpDemuX reverse transcription adapter (RTA) 3, and modification status was identified by construct. Run 2 used WarpDemuX RTAs 3 and 5, one for each modification status, and was demultiplexed with WarpDemuX v1.2.0 (van der Toorn et al., 2025). Run 1 included codon-optimized influenza hemagglutinin (HA) sequences designed with VaxLab (Kim et al., 2026) and synthetic QK sequences; the nine QK constructs together contain 998 of the 1,024 possible 5-mers (to be deposited; see Data availability). QK-03 was excluded from Run 1 because of low quality.

Sequencing data were obtained through MinKNOW and basecalled with Dorado v2.0.0 (Oxford Nanopore Technologies, 2026b). Alignment was done with minimap2 (v2.30, r1287) (Li, 2018, 2021).

### 2.6 Evaluation methods

For the basecaller evaluation (Fig. 2a, b), 12,192 m1Ψ reads were drawn per held-out construct. For the four constructs sequenced in both runs, half were drawn from each run. For the other six constructs, all reads were drawn from Run 2 (121,920 reads; 24,384 from Run 1 and 97,536 from Run 2). Of these, 120,725 reads were assigned to their own construct by both models at seed length k = 7 and window size w = 5, and 118,672 of them had no supplementary alignment under either model. From the public Curlcake dataset, which provides the 1,000 reads with the longest alignments per construct (Cruciani et al., 2025), 1,998 reads of Curlcakes 3 and 4 (999 each) passed the same two filters. Error rate was calculated on these 118,672 and 1,998 reads at novel sites as (substitutions + deletions + insertions) ÷ depth, pooled over reads and positions. Read loss was measured on the 120,725 reads by realigning them with minimap2 at seed lengths k = 7 to 28 (w = 10); a read counts as lost when it is no longer assigned to its own construct.

**Fig. 2.**
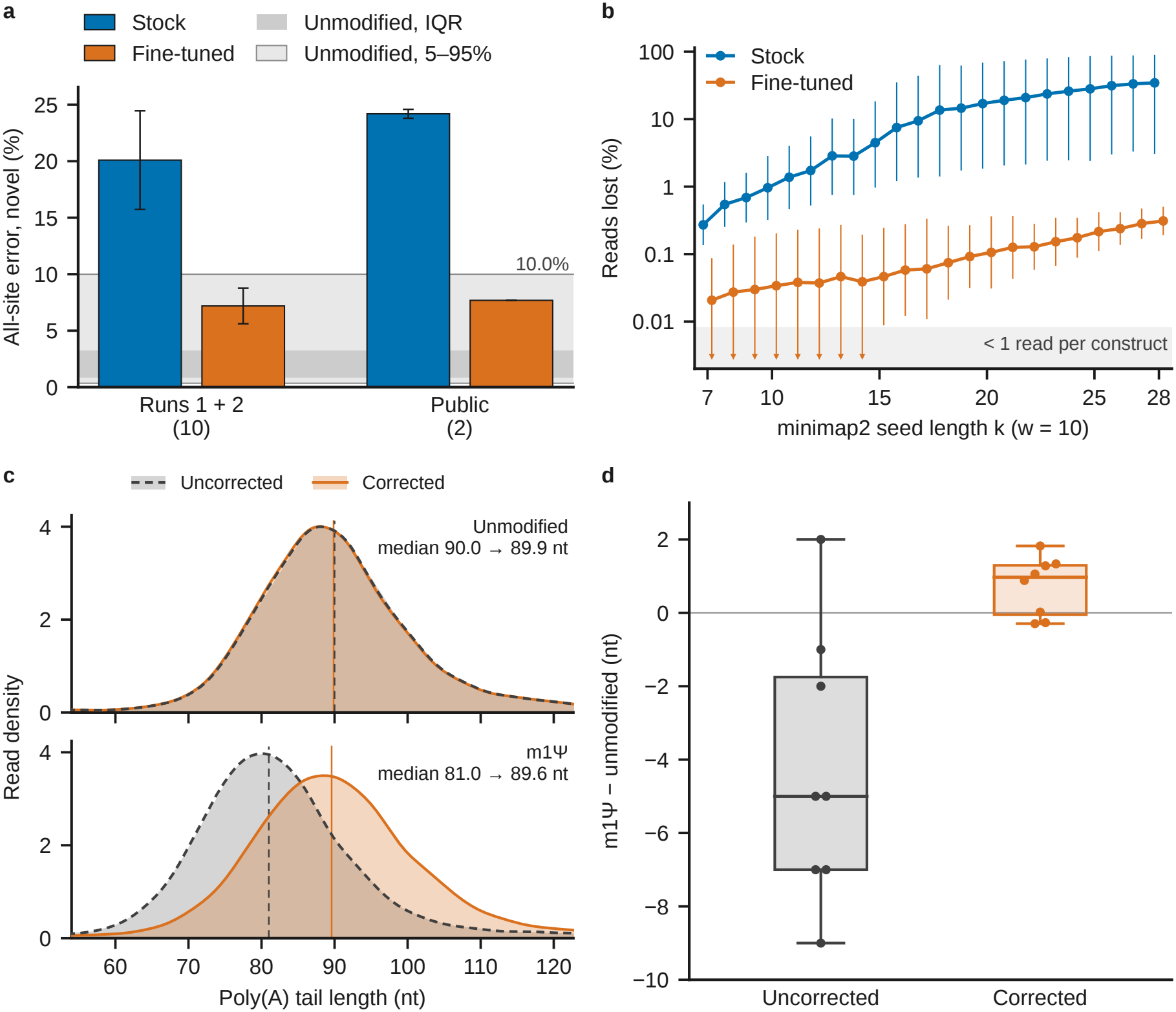
Evaluation of fine-tuned basecaller and poly(A) tail length correction. All panels show constructs that were held out from training; the reads and procedures are described in Sections 2.4 and 2.6. **(a)** Error rate at novel sites of held-out constructs, by basecaller model (118,672 reads for Runs 1 and 2; 1,998 reads for the public dataset). The number below the x-axis is the number of constructs evaluated. Error bars indicate standard deviation (SD) across constructs, weighted by depth. The gray bands show the per-position error rate of unmodified samples in Run 2. The lighter band is the 5%–95% range, and the darker band is the interquartile range (IQR). **(b)** Reads lost during alignment as the minimap2 seed length k increases (w = 10), out of the 120,725 reads that both models assigned to their own construct at k = 7, w = 5. Error bars show the SD across constructs on the logit scale; a bar whose lower end falls below one read per construct is shown as an arrowhead. **(c)** Change in the QK-10 poly(A) tail length distribution upon correction; 11,897 m1Ψ and 10,713 unmodified reads passed the poly(A) read rule. 1-nt bins were smoothed with a Gaussian kernel (2-nt bandwidth). For both modified and unmodified RNA, the solid orange lines mark the median poly(A) tail length after correction, and the gray dashed lines mark the median before correction. **(d)** Difference between the m1Ψ and the unmodified poly(A) median of the same construct, before and after correction, for the eight constructs that have an unmodified arm. The box and its line indicate the IQR and median, and the whiskers indicate the minimum and maximum. Each point represents one construct.

For the poly(A) evaluation (Fig. 2c, d), the m1Ψ and unmodified RNA were both drawn from Run 2, the only run in which constructs were sequenced in both versions, at 12,192 reads per construct per arm, and basecalled with the fine-tuned model. Each construct was corrected with F and E fitted without its own design (leave-one-design-out).

## 3. Usage

### 3.1 Installation and input

mRNAspect runs as a desktop app for macOS on Apple Silicon, or as two command-line tools on Linux and macOS (mrnaspect for finished data, mrnaspect-realtime for a live run). The app carries its own Python environment, including minimap2 v2.30. The Dorado basecall server and the stock basecalling models should be downloaded separately, while the fine-tuned model will be deposited at Zenodo (see Code availability). Installation is described in `README.md` of the GitHub repository (see Code availability). In the app, the path to the Dorado basecall server, the processor, and the directory of basecaller models are set once per device, when launched for the first time. For each run, the user specifies the run mode, input directory, reference sequence, modification status of the sample, basecaller model, and output directory. If the BAM files are selected as input and the read quality filter was applied during basecalling, the user can use only passed reads, or all reads, including failed reads. Construct-specific modification status and parameters for basecalling, alignment, and other QC measurements can be changed through a YAML configuration file.

### 3.2 Demonstration and output

The demo uses data from the first two hours of Run 1, in three forms: the FAST5 file for MinKNOW playback, 40 POD5 files written at 180 s intervals, and BAM files basecalled from those POD5 files with the fine-tuned and stock models. The demo data and the QK reference sequences will be deposited in public repositories (see Data availability). After a simulated MinION device is added in MinKNOW, the FAST5 file can be played back. To simulate real-time analysis, `batch_duration = “180s”` or a shorter value should be set in the default_writer.toml of MinKNOW. A replay can also be simulated with `mrnaspect-realtime --mimic-source-dir`.

Run-level results are displayed once at least 1,000 reads have mapped, and construct-level results are displayed once at least 100 reads have mapped to the construct. Fig. 1b shows mRNAspect running on macOS. The Overview panel of a live mode run displays the run information and the run-level QC results: read count, mapping rate, read length, poly(A) tail length, and read quality. Read count, per-base identity, and full-length fraction are reported per construct. The Overview panel summarizes the run, while details are shown in the other panels and in the reports. The eight panels are Overview, Read length, Poly(A), Quality, Constructs, Identity, Full length, and Sequencing information.

By default, a self-contained HTML interim report is generated each time an input file is processed. It includes both the run-level and the construct-level results. When the run is finished, final reports are generated in HTML and PDF formats. Reports can be accessed through the ‘View interim report’, ‘View PDF report’ (when the run is finished), or ‘Show in Finder’ buttons.

## 4. Evaluation

### 4.1 Performance of fine-tuned model on m1Ψ-modified RNA

We evaluated the performance of the fine-tuned model in Run 1 and Run 2 on the constructs that were held out from training. Fig. 2a shows that fine-tuning reduced the error rate from 20.1% to 7.2% on these constructs, and from 24.2% to 7.7% on the public Curlcake dataset (from 24.5% to 7.7% on Curlcake 4 alone). Relative to unmodified RNA, the m1Ψ error rate fell from 6.5× to 2.4×. By reducing basecalling errors, fine-tuning increased mapping sensitivity across a broad range of minimap2 seed lengths (k). As shown in Fig. 2b, reads basecalled with the fine-tuned model were mostly retained as the seed length increased, whereas reads basecalled with the stock model were increasingly lost. At k = 20, 17.0% of the reads were lost with the stock model, but only 0.1% of the reads were lost with the fine-tuned model, relative to the reads mapped to their own construct at k = 7. Lost reads followed two patterns. At k = 20, in 8 of the 10 constructs the lost reads were shorter than the retained reads. In the other two constructs, lost and retained reads were nearly the same length, yet these constructs lost 50.1% and 86.8% of their reads at k = 20, which altered their fraction of full-length reads. Thus, improved mapping sensitivity can preserve reads for QC and reduce bias in the full-length fraction.

### 4.2 Correction of poly(A) tail length

Fig. 2c shows the poly(A) tail length distribution of one held-out construct (QK-10) before and after correction. Before correction, the m1Ψ median was 81.0 nt, which is 9 nt shorter than the unmodified RNA of the same construct and the designed tail length (90 nt). After correction, the m1Ψ median was 89.6 nt, within 0.3 nt of its unmodified pair (89.9 nt) and 0.4 nt of the designed tail length. Fig. 2d shows the difference between the m1Ψ-modified and unmodified RNA before and after correction for the eight held-out constructs that have an unmodified pair. After correction, the m1Ψ and unmodified medians differed by at most 1.8 nt. This shows that correction is possible when the reference sequence is known.

### 4.3 Real-time analysis

When measured on a single NVIDIA RTX 2080 Ti in a Linux environment, a POD5 file covering 180 s of sequencing (∼5,700 reads) was processed in 38.2 s on average (n = 38, maximum 40.7 s), which is about 21% of the 180 s interval. Basecalling accounted for 80.7% of this time, so the analysis would be slower on a slower GPU. Alignment, QC, and report generation took 7.7%, 10.7%, and 0.6%, respectively. In the demo data with 15 constructs, run-level measurements were available in under 5 min, and per-construct measurements were available for every construct in about 7 min.

## 5. Discussion

mRNAspect provides real-time QC, integrating multiple therapeutic mRNA attributes on the nanopore DRS platform. It connects sequencing with basecalling, alignment, and modularized QC measurements, reducing the time spent waiting for sequencing to finish and for post-hoc analysis. The user gets results sooner through real-time analysis, and with less effort through the GUI. The QC panels in the GUI and the HTML reports show the latest snapshot of the measurements in intuitive plots, so that users can inspect each construct while the run is in progress. For example, a per-position mismatch-rate plot for each construct shows which positions are error-prone, and a heatmap of read start and end positions shows the truncation pattern. The final PDF report includes detailed results at both the run and construct levels and thus serves as a record of the sample’s quality.

Moreover, integration of the fine-tuned basecaller for m1Ψ-modified RNA helps measure sequence identity and alignment with less bias, partly by preserving reads or sequences that would otherwise be dropped during mapping and alignment. The poly(A) correction also reduces the underestimation of poly(A) tail length in m1Ψ-modified RNA. Together, these two features reduce m1Ψ-induced biases and make QC measurements of m1Ψ-modified RNA more interpretable.

The limitation of this study mainly lies in only partially measuring the attributes of therapeutic mRNA, thus not covering double-stranded RNA contamination or 5′ capping. The measurements covered were not compared with orthogonal methods such as liquid chromatography. While the fine-tuned basecaller reduced basecalling error, its error rate still exceeds that of unmodified RNA, and the selection of the learning rate and the epoch was done with the same evaluation result. To address this, the basecaller will be further trained with broader sequence context, and the evaluation will be done with a separate run from training and model selection, to test generalization across runs. The poly(A) tail correction was fitted and evaluated within a single run, with three 3′-end sequences. Also, defective samples were not analyzed in this version of the study; therefore, the sensitivity of the pipeline has yet to be demonstrated. Finally, the app’s real-time throughput has not been demonstrated on macOS, which is the platform of the packaged app. mRNAspect can be further improved by adding more QC measurements and by testing its sensitivity to defective samples and comparing it with orthogonal methods.

mRNAspect aims to serve as a research and characterization tool for rapid assessment of *in vitro* transcribed mRNA, providing both a run-level overview and detailed construct-level results that are updated while the run is ongoing. Looking ahead, mRNAspect may accelerate therapeutic mRNA development and manufacturing by complementing current QC procedures for the attributes it covers.

## Data availability

The demo dataset used in Section 3.2 (the first 2 h of Run 1 as a bulk FAST5 file, 40 POD5 files and unaligned BAM files from both basecalling models, with the reference FASTA and the run configuration) will be deposited at Zenodo in two records, raw signal and basecalled reads. The QK reference sequences will be deposited at figshare; the HA sequences are from Kim et al., 2026. Public data reused in this study: HeLa and HEK293T cDNA-IVT RNA004 reads (Zou et al., 2025; ENA PRJEB80229) and the Curlcake RNA004 PromethION reads (Cruciani et al., 2025; ENA PRJEB82528). The complete raw data for Run 1 and Run 2 and, until the deposits are public, the demo dataset and the QK reference sequences are available from the corresponding author upon reasonable request.

## Code availability

The source code of mRNAspect is available at https://github.com/ChangLabSNU/mRNAspect under the MIT license; the version described here is v0.1.0. mRNAspect runs as a desktop app on macOS (Apple Silicon) and as command-line tools on Linux and macOS. The Dorado basecall server (tested with version 7.13.6) and the stock basecalling models are downloaded separately from Oxford Nanopore Technologies. The fine-tuned basecalling model `m1psi_tuned_v1` will be deposited in its own Zenodo record under the Oxford Nanopore Technologies Public License v1.0, which permits use for research purposes only.

## Author contributions

S.L. conceived the study and performed the cloning, *in vitro* transcription, and nanopore direct RNA sequencing. S.L. developed mRNAspect, fine-tuned the basecaller, analyzed the data, and wrote the manuscript. Contributions of others are listed in the Acknowledgments.

## Acknowledgments

The author thanks Hyeshik Chang for conceiving the study together with the author, for supervising it, and for reviewing the mRNAspect software; Chae Young Kwon for performing the cloning, *in vitro* transcription, and nanopore direct RNA sequencing together with the author; and Dohye Yun for help with *in vitro* transcription.

Generative AI tools (Claude, Anthropic) were used to assist with software development, analysis scripts and language editing; the author reviewed all output and is responsible for the content.

